# Co-translational biogenesis of lipid droplet integral membrane proteins

**DOI:** 10.1101/2021.08.05.455205

**Authors:** Pawel Leznicki, Hayden O. Schneider, Jada V. Harvey, Wei Q. Shi, Stephen High

**Affiliations:** School of Biological Sciences, Faculty of Biology, Medicine and Health, University of Manchester, Manchester, M13 9PT, United Kingdom; Department of Chemistry, Ball State University, Muncie, IN, 47306, USA

**Keywords:** Co-translational, Endoplasmic Reticulum, Lipid Droplets, Membrane Proteins, Protein Targeting

## Abstract

Membrane proteins destined for lipid droplets (LDs), a major intracellular storage site for neutral lipids, are inserted into the endoplasmic reticulum (ER) and then trafficked to LDs where they reside in a hairpin loop conformation. Here, we show that LD membrane proteins can be delivered to the ER either co- or post-translationally and that their membrane-embedded region specifies pathway selection. The co-translational route for LD membrane protein biogenesis is insensitive to a small molecule inhibitor of the Sec61 translocon, Ipomoeassin F, and instead relies on the ER membrane protein complex (EMC) for membrane insertion. Strikingly, this route can also result in a transient exposure of the short N-termini of LD membrane proteins to the ER lumen, followed by topological rearrangements that enable their transmembrane segment to form a hairpin loop and N-termini to face the cytosol. Our study reveals an unexpected complexity to LD membrane protein biogenesis and identifies a role for the EMC during their co-translational insertion into the ER.

**SUMMARY STATEMENT:** Insertion of many lipid droplet membrane proteins into the endoplasmic reticulum (ER) is co-translational, mediated by the ER membrane protein complex (EMC) and involves topology reorientation.

## INTRODUCTION

Lipid droplets (LDs) are highly conserved, dynamic organelles that are present in virtually all eukaryotic cells. They act as the main intracellular storage site for neutral lipids, which can then be used for energy production and membrane biosynthesis. LDs also regulate protein homeostasis (Cermelli et al., 2006; Moldavski et al., 2015) and play a key role in the propagation of certain viruses, including hepatits C (Miyanari et al., 2007; Samsa et al., 2009; Zhang et al., 2018). The massive accumulation of LDs observed in pathological conditions such as obesity, fatty liver disease (steatosis), diabetes and atherosclerosis further underscores their medical relevance whilst mutations in LD proteins are linked to lipodystrophies and motor neuron diseases (Fujimoto and Parton, 2011; Henne et al., 2018; Krahmer et al., 2013).

LD formation is initiated when excess neutral lipids are deposited between the leaflets of the endoplasmic reticulum (ER) bilayer. A growing LD is then formed at discrete ER sites enriched in LD biogenesis factors such as seipin and LDAF1 (Chung et al., 2019). The nascent LD subsequently buds off from the ER into the cytosol (Fujimoto and Parton, 2011; Henne et al., 2018; Krahmer et al., 2013), although there is compelling evidence that LDs can stay connected to the ER via membrane bridges (Wilfling et al., 2014; Wilfling et al., 2013; Zehmer et al., 2009). Hence, LD biogenesis at the ER results in the formation of a hydrophobic oil phase surrounded by a phospholipid monolayer derived from the cytosolic leaflet of the ER. Such an architecture distinguishes LDs from other intracellular membrane-enveloped organelles, which are surrounded by phospholipid bilayers, and has profound consequences for phospholipid packing and protein localisation to LDs (Caillon et al., 2020; Dhiman et al., 2020; Kory et al., 2015; Krahmer et al., 2011; Olarte et al., 2020).

Depending on how they reach their destination, LD-localising proteins can be divided into two distinct classes. Soluble proteins target LDs from the cytosol, using amphipathic helices, lipid anchors and protein-protein interactions to associate with LDs, and their binding is defined by protein crowding and phospholipid packing defects within the LD-surrounding monolayer (Chorlay and Thiam, 2020; Dhiman et al., 2020; Kory et al., 2015). In contrast, integral membrane proteins must first be inserted into the ER membrane prior to their trafficking to LDs (Ingelmo-Torres et al., 2009; Ostermeyer et al., 2004; Schrul and Kopito, 2016; Stevanovic and Thiele, 2013; Turro et al., 2006; Zehmer et al., 2009; Zehmer et al., 2008). Such integral membrane proteins include a number of lipid synthesising and metabolising enzymes which relocate from the ER to LDs to support LD growth (Wilfling et al., 2013), limit their lipolysis (Olzmann et al., 2013; Zhang et al., 2017) and potentially link LDs to protein quality control machinery (Klemm et al., 2011; Spandl et al., 2011). The energetic barrier associated with accommodating hydrophilic residues within the very hydrophobic LD interior, combined with the reduced thickness of the LD phospholipid monolayer as compared to phospholipid bilayers, mean that LD integral membrane proteins adopt a hairpin or amphipathic helix conformation where both their N- and C-termini face the cytosol (Caillon et al., 2020; Olarte et al., 2020; Thiam et al., 2013). Indeed, fully membrane spanning proteins are specifically excluded from LDs (Khaddaj et al., 2022).

The majority of integral membrane proteins destined for the secretory pathway are co-translationally targeted to, and inserted into, the ER membrane via a pathway that is initiated by the signal recognition particle (SRP) (O’Keefe et al., 2021a). Hence, SRP binds to a hydrophobic N-terminal signal sequence or the first transmembrane domain (TMD) of a nascent membrane protein as soon as it emerges from the ribosome, and targets the ribosome-nascent chain complex to the ER where SRP interacts with its membrane-tethered receptor, SR. In most cases, the ribosome-nascent chain complex is then transferred to the Sec61 translocon, composed of Sec61α, Sec61β and Sec61γ, which opens laterally into the ER membrane to enable the release of newly synthesised TMD(s) into the bilayer (O’Keefe et al., 2021a). One hallmark of this Sec61-dependent pathway is its sensitivity to small molecules, including Ipomoeassin F, which strongly inhibit membrane protein integration at the ER (Luesch and Paavilainen, 2020; McKenna et al., 2017; O’Keefe et al., 2021b; Zong et al., 2019). Indeed, the insensitivity of so-called type III membrane proteins, defined by the lack of an N-terminal cleavable signal sequence and translocation of the N-terminus to the ER lumen, to these small molecule inhibitors (McKenna et al., 2017; Zong et al., 2019) led to the identification of an alternative pathway for their co-translational integration into the ER membrane (O’Keefe et al., 2021c). This alternative pathway for the co-translational integration of type III membrane proteins relies on the ER membrane complex (EMC) (O’Keefe et al., 2021c), consistent with growing evidence that it can act as a stand-alone membrane insertase for certain types of TMDs (Bai et al., 2020; Chitwood et al., 2018; Guna et al., 2018; O’Donnell et al., 2020; Pleiner et al., 2020).

Relatively little is known about the biogenesis of LD membrane proteins at the ER. Studies of oleosins, major components of plant LDs, established that, at least in heterologous plant/mammalian and plant/yeast systems (Abell et al., 2002; Beaudoin et al., 2000), these proteins are targeted to the ER co-translationally via the action of SRP and SR. However, oleosins are characterised by a very hydrophobic membrane-embedded region of ∼72 aa (Murphy, 1993), which is substantially longer than the ∼30 aa hairpin found in most LD membrane proteins, and hence the generality of these findings has been questioned (Dhiman et al., 2020). More recently, Schrul and Kopito (Schrul and Kopito, 2016) carried out an elegant analysis of the ER membrane insertion requirements for a model LD membrane protein, UBXD8. They found that UBXD8 is post-translationally inserted into the ER membrane with a preformed hairpin loop topology via a process that is mediated by soluble Pex19 and its membrane-tethered receptor, Pex3 (Schrul and Kopito, 2016).

Likewise, two reticulon-homology domain containing proteins, Arl6IP1 and Rtn4C, that reside in the ER membrane in a hairpin loop conformation are also suggested to follow this post-translational pathway (Yamamoto and Sakisaka, 2018). However, the biogenesis of other LD integral membrane proteins has not been studied in detail leaving open the possibility of alternative biosynthetic pathways, as previously identified for tail-anchored membrane proteins (Casson et al., 2017).

In order to better understand their biogenesis at the ER, we have used a panel of well-defined LD membrane proteins and investigated the requirements for their delivery to, and insertion into, the ER membrane. We find that LD membrane proteins rely on at least two distinct biosynthetic pathways which are selected via the hydrophobic, membrane-inserting, region of individual LD proteins. One is a post-translational route that is used by UBXD8 and HSD17B7 and appears equivalent to the previously defined Pex19/Pex3-mediated pathway (Schrul and Kopito, 2016). The other is a co-translational route that is favoured by the majority of the LD membrane proteins we have tested. This co-translational pathway is mediated by the EMC and involves a transient exposure of the short N-termini of LD membrane proteins to the ER lumen prior to their assembly into nascent LDs.

## RESULTS

### LD membrane proteins differ in their requirements for insertion into the ER

To investigate whether LD membrane proteins share a common pathway for their biogenesis at the ER, we compared the ER membrane insertion requirements of the well-studied UBXD8 with those of other known LD membrane proteins (Fig. 1A). Initially, we adopted the homologous *in vitro* translation system that was instrumental in identifying the Pex19/Pex3-dependent route for UBXD8 biogenesis (Schrul and Kopito, 2016). Furthermore, by carrying out LD membrane protein synthesis either in the presence of ER-derived membrane (co-translationally), or by adding the ER membrane following translation termination (post-translationally), we were able to establish the favoured mode of membrane binding/insertion for each LD protein studied (McKenna et al., 2016; Schrul and Kopito, 2016).

**Fig. 1.**
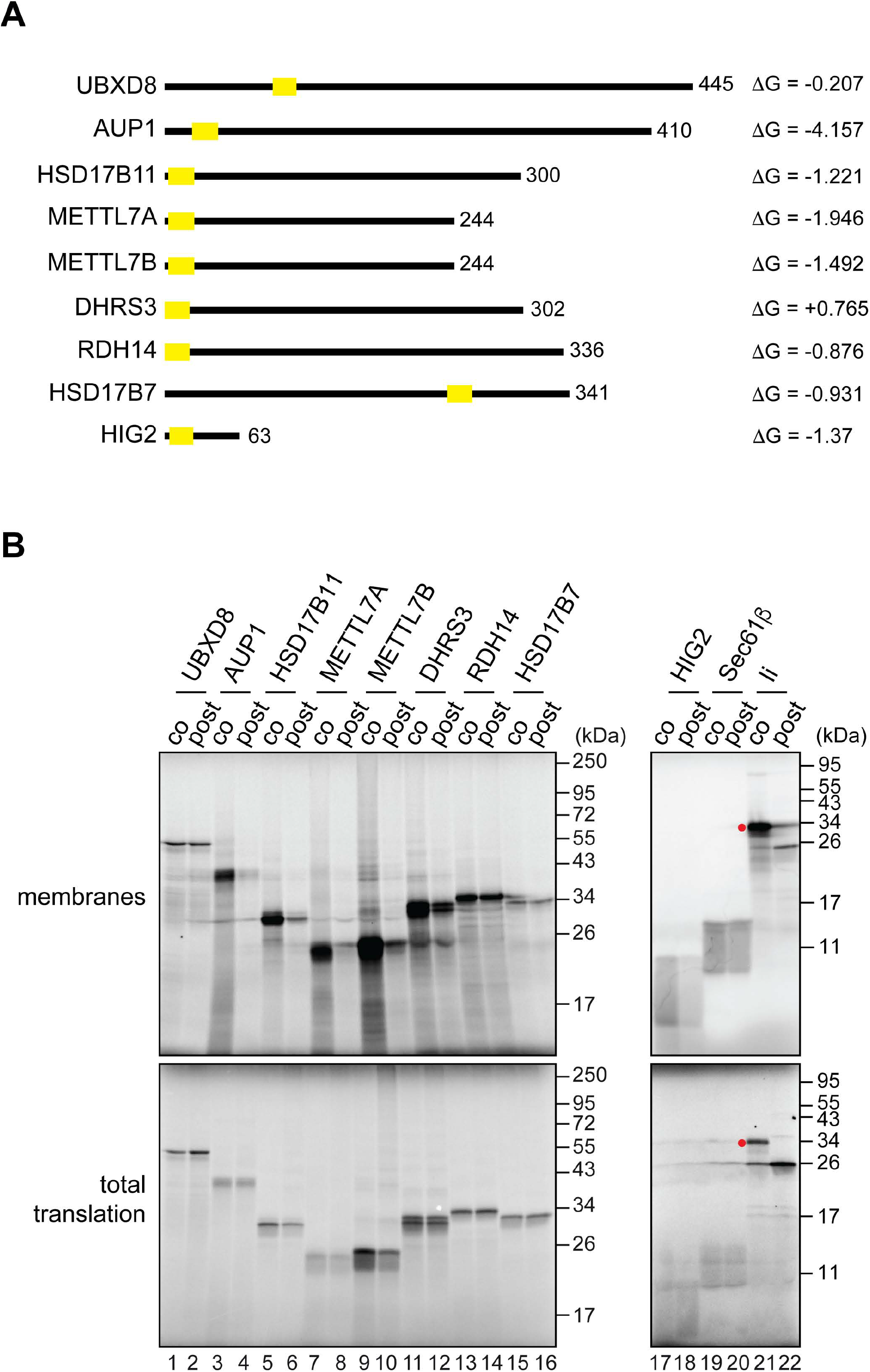
LD membrane proteins differ in their requirements for delivery to the ER. **(A)** A schematic representation of the LD membrane proteins used in this study. Protein length, localisation of predicted TMDs (yellow rectangles) (ΔG predictor, (Hessa et al., 2007)) and estimated ΔG (in kcal/mol) associated with their insertion into the ER membrane (Hessa et al., 2007) are indicated. **(B)** Untagged membrane proteins as indicated were synthesised *in vitro* using rabbit reticulocyte lysate and ER-derived microsomes, which were either present throughout the reaction (co-translational conditions; “co”) or added following translation termination (post-translational conditions; “post”). Membrane-associated material (top panels) and total translation reactions (bottom panels) were resolved by SDS-PAGE, and results were visualised by phosphorimaging. Sec61β and invariant chain (Ii; also known as HLA class II histocompatibility antigen gamma chain/CD74) are control proteins inserted into the ER membrane either post- or co-translationally, respectively. Red dot indicates the N-glycosylated species of Ii.

As previously reported (Schrul and Kopito, 2016), we found that UBXD8 bound to ER-derived microsomes equally well under co- and post-translational conditions akin to Sec61β, a member of so-called tail-anchored membrane proteins that are delivered to the ER post-translationally (O’Keefe et al., 2021a) (Fig. 1B, lanes 1 and 2, 19 and 20). Furthermore, this behaviour was mirrored at a qualitative level by two other LD membrane proteins, RDH14 and HSD17B7 (Fig. 1B, lanes 13 and 14, 15 and 16). However, in contrast to these three examples, the membrane association of the majority of the other LD proteins analysed was dramatically reduced under conditions that require post-translational targeting to the ER (Fig. 1B, lanes 3-10, 17 and 18), despite comparable levels of protein synthesis (Fig. 1B, bottom panel). Hence, these LD proteins behaved much like the invariant chain (Ii, also known as HLA class II histocompatibility antigen gamma chain/CD74) (Fig. 1B, cf. lanes 21 and 22), a well-studied substrate for the Sec61-mediated, co-translational pathway of membrane insertion into the ER (Lipp and Dobberstein, 1986; Zong et al., 2020; Zong et al., 2019). In the case of DHRS3 we observed an intermediate effect, as evidenced by a modest reduction in its membrane association under post-translational conditions (Fig. 1B, cf. lanes 11 and 12). We note that the TMD of DHRS3 appears less hydrophobic than any of the other LD membrane proteins studied (see Fig. 1A) and hence our data are consistent with the possibility that DHRS3 associates with the ER via an amphipathic helix rather than a fully membrane inserted hairpin loop (Pataki et al., 2018).

Taken together, we conclude that LD membrane proteins differ in their capacity to be post-translationally inserted into the ER. Unlike UBXD8, it may be that other LD proteins are not effectively maintained in a membrane insertion competent form by cytosolic chaperones such as Pex19 (Schrul and Kopito, 2016) or, alternatively, they employ a co-translational pathway for membrane insertion.

### LD membrane proteins expose their N-termini to the ER lumen

Whilst the TMD of UBXD8 has been proposed to insert post-translationally into the ER membrane as a hairpin loop (Schrul and Kopito, 2016), the majority of membrane proteins that are co-translationally inserted into the ER expose hydrophilic regions that flank their TMD to the ER lumen (O’Keefe et al., 2021a). Given our finding that LD membrane proteins may not follow a single biosynthetic pathway, we wondered whether some LD membrane proteins may also translocate their short hydrophilic N-terminus into the ER lumen during biogenesis. To address this question, we tagged our panel of LD membrane proteins (see Fig. 1A) with a short N-terminal extension derived from bovine rhodopsin (OPG2), which contains two N-glycosylation sites (Leznicki et al., 2010; McKenna et al., 2016; O’Keefe et al., 2021b; O’Keefe et al., 2021c; Pedrazzini et al., 2000; Schrul and Kopito, 2016) and with a C-terminal FLAG epitope. As N-glycosylation is an ER lumen-specific modification, the presence of N-glycan(s) on the OPG2 tag can be used as a read-out for its translocation to the ER lumen (Leznicki et al., 2010; McKenna et al., 2016; O’Keefe et al., 2021b; O’Keefe et al., 2021c; Pedrazzini et al., 2000; Schrul and Kopito, 2016).

Strikingly, when these OPG2-tagged LD membrane proteins were synthesised *in vitro* in the presence of ER-derived microsomes most of them migrated as two distinct species (Fig. 2A for membrane fraction and Fig. S1 for total translation products). By means of their sensitivity to Endoglycosidase H (EndoH), we established that in each case the higher molecular weight species were N-glycosylated (Figs 2A and S1, red dots) and hence their OPG2 tag had entered the ER lumen. The N-terminal location of the predicted TMD for most of the proteins analysed (see Fig. 1A) means that the distance between the N-glycosylation sites of the OPG2 tag and the active site of the oligosaccharyltransferase complex is relatively short (cf. (Nilsson and von Heijne, 1993)). Hence, only the distal N-glycosylation site of the OPG2 tag is modified (cf. (Nilsson and von Heijne, 1993)). Only three of the OPG2-tagged LD proteins showed no evidence of N-glycosylation: UBXD8, DHRS3 and HSD17B7 (Fig. 2A, cf. lanes 1 and 2, 11 and 12, 15 and 16), consistent with previous studies of UBXD8 (Schrul and Kopito, 2016) and DHRS3 (Pataki et al., 2018). In short, we find that the majority of LD proteins which preferentially associate with the ER membrane under co-translational conditions (cf. Fig. 1B), can also be N-glycosylated when tagged at their N-terminus with an OPG2 extension (Fig. 2A). This behaviour supports a model where this group of LD membrane proteins share a common, most likely co-translational, pathway for their biogenesis at the ER. Importantly, the OPG2 and FLAG tags do not alter the biosynthetic pathway selection of the LD proteins tested (Fig. S2), nor do they affect their delivery to LDs in oleic acid-loaded U2OS cells (Fig. S3).

**Fig. 2.**
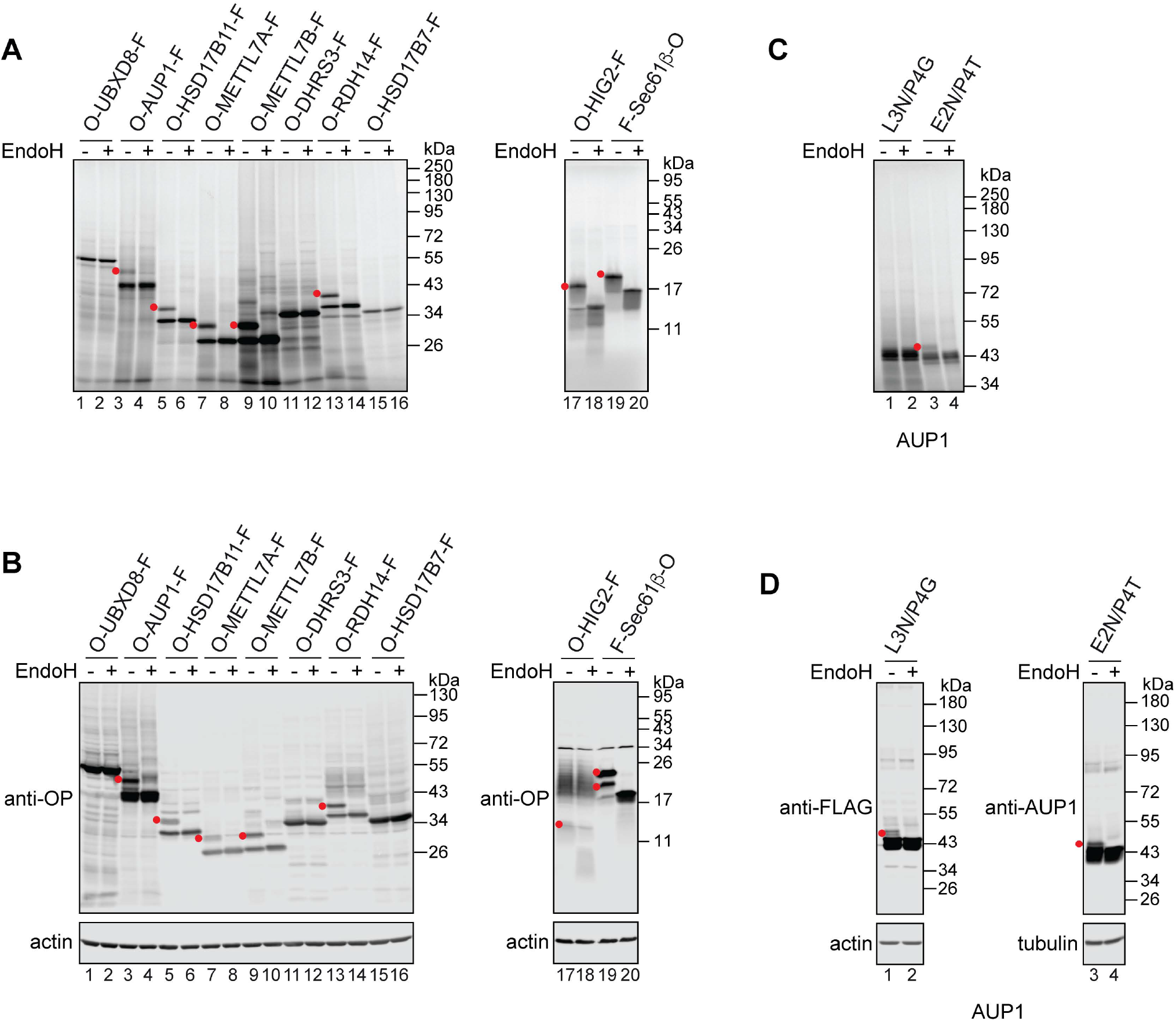
LD membrane proteins translocate their N-termini to the ER lumen. **(A)** Indicated LD membrane proteins bearing an N-terminal rhodopsin-derived (OPG2) tag (“O”), which contains two N-glycosylation sites, and a C-terminal FLAG epitope (“F”) were translated *in vitro* in the presence of ER-derived microsomes. Membranes were isolated, N-glycosylated species (red dots) identified based on their altered electrophoretic mobility in SDS-PAGE following Endoglycosidase H (EndoH) digestion, and results visualised by phosphorimaging. Sec61β with an N-terminal FLAG and a C-terminal OPG2 tags was used as an ER-resident control protein which fully spans the lipid bilayer. **(B)** Proteins used in (A) were transiently expressed in U2OS cells, which were then lysed and, where indicated, treated with EndoH. Samples were resolved by SDS-PAGE and results visualised by Western blotting using anti-rhodopsin (OP) and anti-actin antibodies. **(C**,**D)** Indicated variants of AUP1 with (AUP1^L3N/P4G^) or without (AUP1^E2N/P4T^) a C-terminal FLAG epitope were translated *in vitro* in the presence of ER-derived microsomes (panel C) or transiently expressed in U2OS cells (panel D), and their N-glycosylation status was tested by means of EndoH sensitivity. Results were visualised either by phosphorimaging (panel C) or by Western blotting using the indicated antibodies (panel D).

To test our *in vitro* findings in a cell-based system, we transiently expressed the same tagged LD membrane proteins in U2OS cells and used EndoH sensitivity to test their N-glycosylation (Fig. 2B). We obtained qualitatively similar results to the *in vitro* assay (cf. Figs 2A and 2B), supporting the physiological significance of our findings. Since the N-terminus of neither UBXD8 nor HSD17B7 reaches the ER lumen (Figs 2A and 2B, cf. lanes 1 and 2, 15 and 16) we wondered if they may assume an opposite topology and translocate their C-termini across the ER membrane and to address this question, we reversed the localisation of the OPG2 and FLAG epitopes. Since we could not detect any N-glycosylation of the OPG2 tag when placed at either the N- or C-terminus of these two proteins, either *in vitro* (Fig. S4A) or in cultured cells (Fig. S4B), we conclude that UBXD8 and HSD17B7 follow a distinct biosynthetic pathway from the majority of LD membrane proteins we have tested. This pathway is characterised by its efficient operation under post-translational conditions and the lack of access to the ER lumen that it provides LD proteins at the ER membrane (cf. Figs 1B, 2A, 2B and S4). Only in the case of RDH14 did we observe comparable levels of membrane association under co- and post-translational conditions and the efficient N-glycosylation of its N-terminal OPG2 tag (Figs. 1B, 2A and 2B). We conclude that RDH14 can likely access both co- and post-translational routes for LD membrane protein biogenesis at the ER (see Discussion).

Finally, to test whether we can detect LD membrane protein exposure to the ER lumen in the absence of an artificial OPG2 tag we took advantage of the fact that the predicted TMD of AUP1 results in a slightly longer N-terminal region than in our other “co-translational” LD membrane proteins (see Fig. 1A). This allowed us to create two AUP1 variants, AUP1^L3N/P4G^ and AUP1^E2N/P4T^, that bear single N-glycosylation sites engineered into their otherwise native sequence. We found that a fraction of AUP1^E2N/P4T^ was N-glycosylated both *in vitro* (Fig. 2C, cf. lanes 3 and 4, and Fig. S1) and in cells (Fig. 2D, cf. lanes 3 and 4) whilst AUP1^L3N/P4G^ was N-glycosylated in cells, but not *in vitro* (Figs. 2C and 2D cf. lanes 1 and 2, and Fig. S1). Interestingly, a previous study of AUP1 (Stevanovic and Thiele, 2013) failed to detect N-glycosylation of the AUP1^L3N/P4G^ variant in COS-7 cells indicating that modification of this particular engineered site may be sensitive to the experimental system used, as further evidenced by the difference between the *in vitro* and cell-based assays we observe with this variant. Taken together, our results strongly suggest that during their biogenesis at the ER a number of LD membrane proteins can translocate their short N-terminal hydrophilic domain into the ER lumen.

### Co-translationally inserted LD membrane proteins access the ER lumen transiently before being trafficked to LDs

The N-glycosylation of LD membrane proteins that we detected was typically incomplete (see Fig. 2) raising the possibility that LD membrane proteins with their N-termini translocated into the ER lumen may represent a distinct population of polypeptides that stably reside in the ER. Since N-glycosylation occurs specifically in the ER lumen we reasoned that the presence of an N-glycosylated LD membrane protein on LDs would confirm that having accessed the ER lumen, such proteins remain competent for sorting into LDs. Hence, we used flotation through a sucrose density gradient to isolate LDs (Ingelmo-Torres et al., 2009) from oleic acid-loaded HepG2 cells that were transiently expressing OPG2-HSD17B11-FLAG.

A clear separation of LDs from the ER was achieved with LDs floating to fraction 1 at the top of the gradient (Fig. 3A, lane 6), while the ER being distributed more broadly with the bulk of it located in fractions 3, 4 and 5 (see Fig. 3A, lanes 2, 3 and 4). The N-glycosylated form of OPG2-HSD17B11-FLAG was located in two distinct fractions, with the strongest signal recovered in the LD containing fraction 1 (Fig. 3A, lane 6, “1g”), together with most of non-glycosylated OPG2-HSD17B11-FLAG. The second pool of N-glycosylated OPG2-HSD17B11-FLAG was found in the middle of the ER-containing fractions (Fig. 3A, lane 3, “1g”). In order to establish that the N-glycosylated OPG2-HSD17B11-FLAG recovered in the LD-enriched fraction 1 is authentically LD-associated we calculated the potential ER contamination of this fraction, relative to the peak ER-containing fraction 4, by quantifying their respective levels of BAP31 (see “Materials and Methods” section). Using this criterion we conclude that the vast majority of N-glycosylated OPG2-HSD17B11-FLAG (∼89%) is authentically bound to LDs, together with a substantial fraction of the non-glycosylated protein. Hence, only ∼11% of the N-glycosylated OPG2-HSD17B11-FLAG recovered in the LD fraction can be ascribed to contamination arising from its ER-localised form (Fig. 3B).

**Fig. 3.**
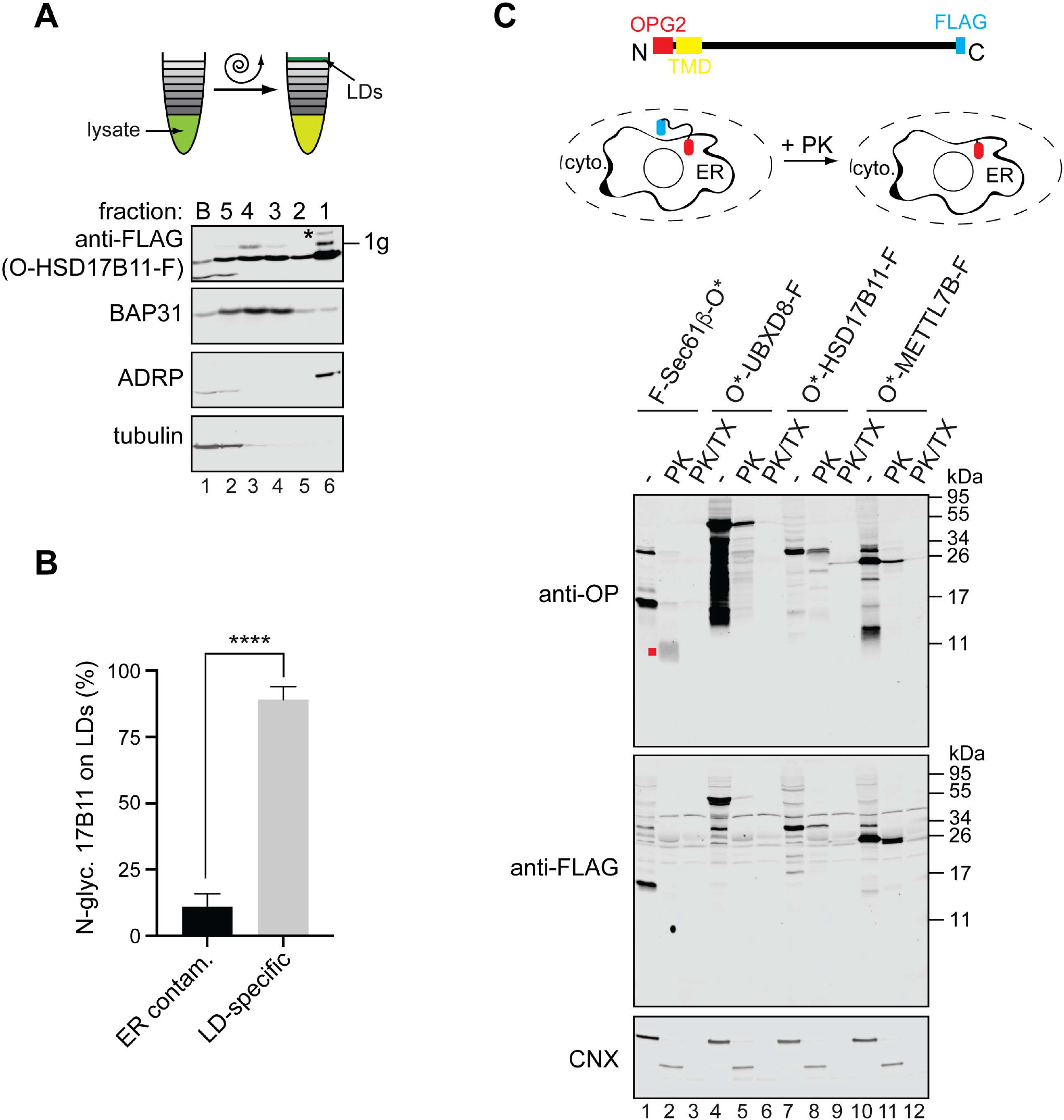
LD membrane proteins co-translationally inserted into the ER reach LDs. **(A)** HSD17B11 with an N-terminal OPG2 tag and a C-terminal FLAG epitope (O-HSD17B11-F) was expressed in HepG2 cells, which were subsequently loaded with oleic acid in the presence of 50 μM zVAD-fmk to inhibit cytosolic N-glycanase (Misaghi et al., 2004). Cells were homogenised, LDs isolated from the post-nuclear supernatant by flotation through a sucrose gradient, and fractions analysed for the presence of selected intracellular compartments by Western blotting with organelle-specific markers: BAP31 (ER), ADRP (LDs) and tubulin (cytosol). Migration of ectopically expressed OPG2-HSD17B11-FLAG was established by Western blotting with the anti-FLAG antibody. “1g” indicates singly N-glycosylated OPG2-HSD17B11-FLAG whilst “*” denotes an EndoH-resistant species most likely cross-reacting with the anti-FLAG antibody. **(B)** The proportion of N-glycosylated OPG2-HSD17B11-FLAG in the LD fraction (see panel A) that corresponds to contaminating ER membranes and to authentic LD-associated species was calculated based on the relative levels of an ER marker, BAP31, in each fraction (see Materials and Methods). Quantifications are given as means with error bars indicating standard error of means. Statistical significance was calculated using unpaired t test for n = 4 biological replicates. **** p<0.0001. **(C)** A schematic representation of the ectopically expressed LD membrane proteins and experimental setup used to address their accessibility to proteinase K (PK). Indicated LD membrane proteins and a control, fully membrane-spanning protein, Sec61β, tagged with the FLAG epitope (“F”) and a variant of the OPG2 tag with mutated N-glycan consensus sites (“O*”), were transiently expressed in HeLa cells. The plasma membrane was then selectively permeabilised and protease accessibility tested either in the absence or presence of 1% (v/v) Triton X-100 (TX). Samples were resolved by SDS-PAGE and results visualised by Western blotting with anti-rhodopsin (OP) and anti-FLAG antibodies. Western blotting for an endogenous ER membrane protein, calnexin (CNX), was used to compare ER membrane integrity between samples and provide an internal control. Filled red square indicates a membrane-protected, protease-resistant fragment.

Neither charged amino acids, nor hydrophilic sugar moieties, are readily accommodated by the hydrophobic interior of LDs (Thiam et al., 2013) and the LD phospholipid monolayer is incompatible with the extended conformation of a “classical” helical transmembrane span (Khaddaj et al., 2022; Thiam et al., 2013). We therefore speculate that both the non-glycosylated and N-glycosylated forms of OPG2-HSD17B11-FLAG must assume a hairpin topology in the LD membrane. In the case of the N-glycosylated form of OPG2-HSD17B11-FLAG where we know its N-terminus has been exposed to the ER lumen, we postulate that its topology must at some stage have been rearranged from an ER bilayer-spanning integral TMD to a LD monolayer-inserted hairpin loop arrangement.

To further test this hypothesis, we carried out a protease protection assay on plasma membrane-permeabilised HeLa cells expressing LD membrane proteins tagged with an OPG2 tag at their N-terminus and a FLAG tag at their C-terminus (Fig. 3C). Since N-glycosylation can influence the transmembrane orientation of proteins at the ER (Goder et al., 1999), we used a modified variant of the OPG2 tag which is incapable of being N-glycosylated. In parallel, we analysed the protease accessibility of a model ER-resident membrane protein, Sec61β, tagged with the same epitopes albeit at opposite termini to reflect its membrane topology. We found that proteinase K digested epitope tagged Sec61β generates a membrane-protected, ER lumenal fragment (Fig. 3C, cf. lanes 1 and 2, “anti-OP” panel) which was lost following disruption of the ER membrane with Triton X-100 (Fig. 3C, cf. lanes 1-3, “anti-OP” panel). In contrast, although in some cases a fraction of the full-length protein was refractive to any digestion in the absence of detergent, no comparable membrane protected fragments were apparent for any of the LD membrane proteins tested (Fig. 3C, cf. lanes 4-12, “anti-OP” panel). This behaviour included METTL7B and HSD17B11 that had displayed efficient N-glycosylation at the N-terminus in previous assays (cf. Figs 2A, 2B and 3A). This suggests that whilst the N-termini of these proteins are transiently exposed to the ER lumen, where they become accessible to the N-glycosylation machinery, they do not normally reside in a stable, membrane-spanning, topology at the ER.

In further support of a model where a subset of LD membrane proteins can transiently span the ER membrane during their biogenesis, we found that N-glycosylation of METTL7B and HSD17B11 bearing a wild-type OPG2 tag effectively “traps” a fraction of these proteins in a fully membrane-spanning topology. This N-glycan dependent trapping results in the presence of protease-protected, EndoH-sensitive, species that are absent without such N-glycosylation (cf. Fig. S5 and Fig. 3C, “anti-OP” panels). In contrast, the FLAG epitope placed at the C-terminus of the LD membrane proteins was consistently accessible to proteinase K confirming its cytosolic localisation (Figs 3C and S5, “anti-FLAG” panels). On the basis of these data, we conclude that the N-termini of newly synthesised LD membrane proteins which follow the co-translational route access the ER lumen only transiently and do not stably reside in a membrane-spanning orientation (see “Discussion”).

### Membrane-embedded region specifies pathway selection

We next asked what determines LD membrane protein entry into either the co- or post-translational biosynthetic pathway at the ER (cf. Figs 1 and 2). We speculated that this choice might depend on the properties of the hydrophobic region, which mediates targeting to and insertion into the ER, and/or the presence of a longer hydrophilic domain on the N-terminal side of this region as observed for both UBXD8 and HSD17B7 (see Fig. 1A). To test this hypothesis, we generated chimaeras between METTL7B and UBXD8, two LD membrane proteins that take distinct routes to the ER (cf. Figs 1B, 2A and 2B). Hence, we replaced the predicted TMD of METTL7B with the hairpin motif of UBXD8 (Fig. 4A, METTL7B^UBXD8-TMD^). In parallel, we substituted UBXD8 hairpin loop with the predicted TMD of METTL7B (UBXD8^L7B-shortTMD^) or with the slightly longer region experimentally shown to mediate both ER membrane insertion and efficient trafficking to LDs (UBXD8^L7B-longTMD^ (Zehmer et al., 2008), see legend to Fig. 4A for details of the constructs used). We then assessed the N-glycosylation of the N-terminal OPG2 tag for each of these constructs.

**Fig. 4.**
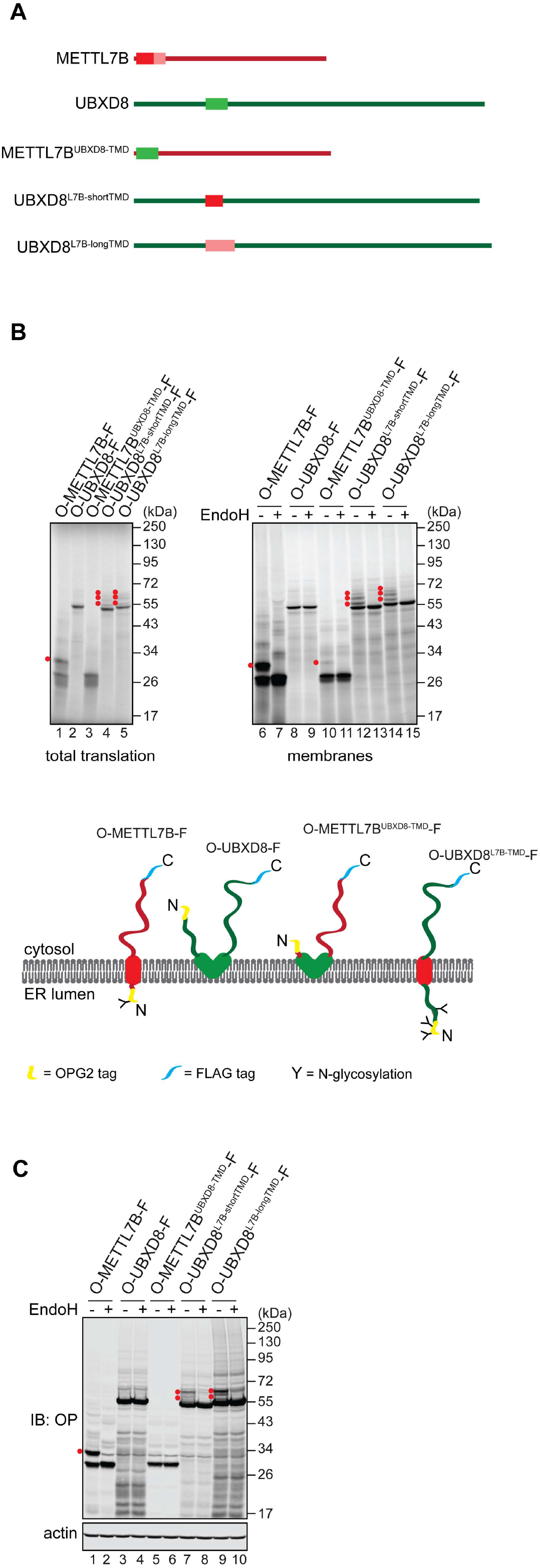
The membrane-embedded region of LD membrane proteins specifies pathway selection. **(A)** A schematic representation of the constructs used in Fig. 4. METTL7B^UBXD8-TMD^ – METTL7B with its predicted TMD (residues 3-26, (Hessa et al., 2007)) replaced with a putative hairpin of UBXD8 (residues 91-119, (Zehmer et al., 2009)), UBXD8^L7B-shortTMD^ – UBXD8 with its hairpin region replaced with residues 3-26 of METTL7B, UBXD8^L7B-longTMD^ – UBXD8 with its hairpin region replaced with residues 3-40 of METTL7B. **(B)** METTL7B and UBXD8 variants (see panel A), tagged with an N-terminal OPG2 (“O”) and a C-terminal FLAG (“F”) epitopes, were translated *in vitro* in rabbit reticulocyte lysate in the presence of ER-derived microsomes. Membranes were isolated and, where indicated, treated with EndoH, followed by SDS-PAGE and phosphorimaging. Red dots indicate N-glycosylated protein species. Schematic representation of each chimeric protein topology based on our experimental results is also shown using the same colour coding as in panel A. **(C)** Protein variants used in (B) were transiently expressed in U2OS cells, which were then lysed and, where indicated, treated with EndoH. Samples were resolved by SDS-PAGE and results visualised by Western blotting with antibodies against rhodopsin (OP) and actin.

As previously, OPG2-METTL7B-FLAG was efficiently N-glycosylated whilst N-glycosylation of OPG2-UBXD8-FLAG was undetectable both *in vitro* and in cells (Fig. 4B, cf. lanes 6-9, and Fig. 4C, cf. lanes 1-4). Strikingly, N-glycosylation of OPG2-tagged METTL7B was virtually absent when its predicted TMD was replaced with the hairpin motif taken from UBXD8 (Fig. 4B, cf. lanes 10 and 11, and Fig. 4C, cf. lanes 5 and 6). Conversely, substituting the UBXD8 hairpin with either of the two versions of the hydrophobic region from METTL7B resulted in the clear N-glycosylation of these UBXD8 variants (Fig. 4B, cf. lanes 12-15, and Fig. 4C, cf. lanes 7-10). Interestingly, in the cell-free system we detected a faint band that corresponds to a triply N-glycosylated form of OPG2-UBXD8^L7B-shortTMD^-FLAG and OPG2-UBXD8^L7B-longTMD^-FLAG despite the OPG2 tag having just two N-glycosylation sites (Fig. 4B, lanes 12-15). This most likely reflects the inefficient N-glycosylation of an endogenous, non-consensus, Asn residue located in the relatively long N-terminal region of UBXD8, that is translocated to the ER lumen in the chimaeras used (Breitling and Aebi, 2013; Dutta et al., 2017). Taken together, we conclude that biosynthetic pathway selection is primarily determined by the membrane-inserting region of LD membrane proteins.

### Co-translational insertion of LD membrane proteins relies on the EMC

In order to investigate the molecular basis for co-translational insertion of LD membrane proteins into the ER membrane, we took advantage of the small molecule Ipomoeassin F (Ipom-F) which selectively inhibits the Sec61-mediated integration of membrane proteins at the ER (O’Keefe et al., 2021b; O’Keefe et al., 2021c; Zong et al., 2020; Zong et al., 2019). When an *in vitro* membrane insertion assay of OPG2-tagged LD membrane proteins was performed in the presence of Ipom-F, their membrane insertion, as judged by the N-glycosylation of their N-terminal region, was unaffected (see Fig. 5A for topology of the proteins used and Figs 5B and 5C, lanes 1-10). In contrast, the N-glycosylation of a classical Sec61 substrate, the type II membrane protein Ii, was almost completely blocked (Figs 5B and 5C, cf. lanes 13 and 14) in line with previous studies (Zong et al., 2020; Zong et al., 2019). Hence, the behaviour of these LD proteins mirrors that of clients that do not require the insertase activity of the Sec61 complex for their integration (Figs 5B and 5C, cf. lanes 11 and 12, 15 and 16), namely Vpu, a type III membrane protein which utilises the EMC (O’Keefe et al., 2021c) and Sec61β, a tail-anchored protein which can access multiple pathways for membrane insertion at the ER (Casson et al., 2017; Guna et al., 2018). On this basis, we concluded that the translocation of the N-termini of LD membrane proteins into the ER lumen does not require translocation via the Sec61 complex.

**Fig. 5.**
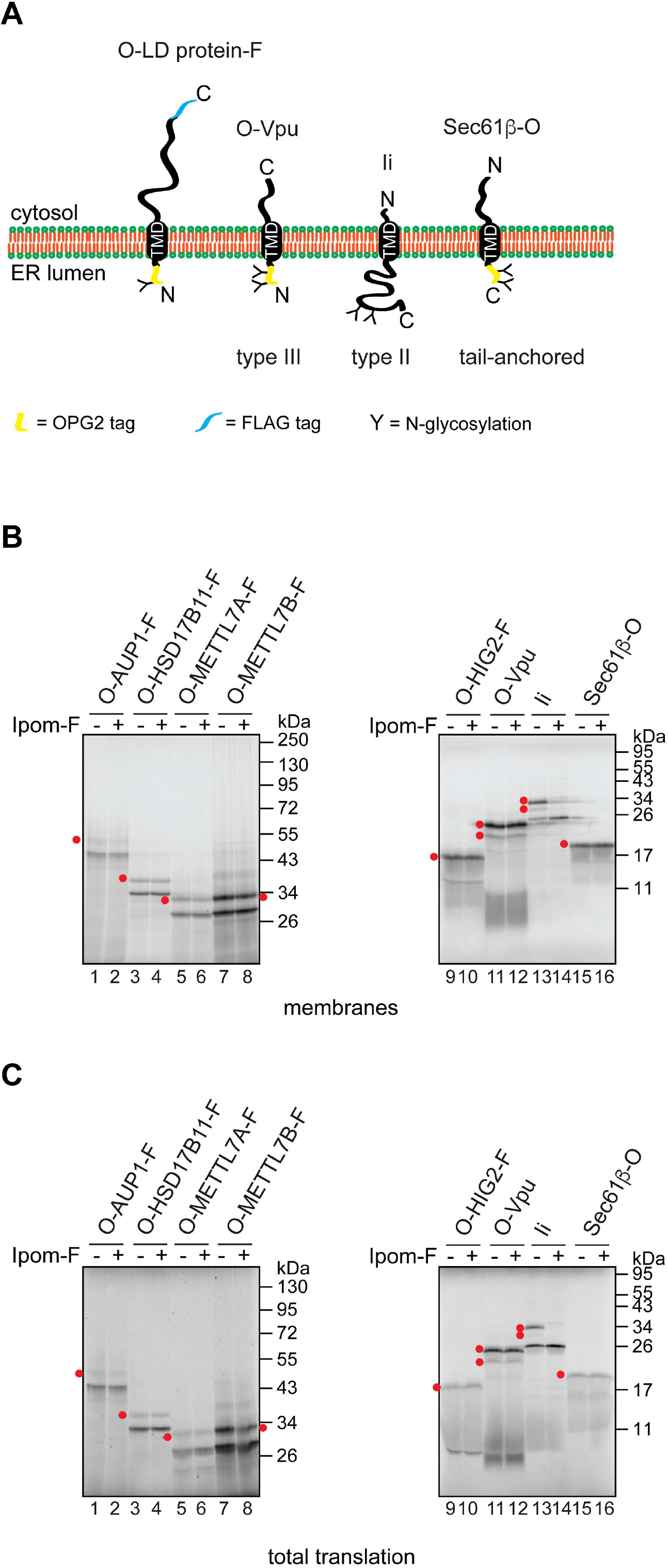
Insertion of LD membrane proteins into the ER is insensitive to Ipomoeassin F. **(A)** Topology of the membrane proteins used in Fig. 5. “O” and “F” indicate the OPG2 and FLAG tags, respectively, whilst “TMD” corresponds to transmembrane domain. **(B**,**C)** Indicated membrane proteins were translated *in vitro* in the presence of ER-derived microsomes and 1 μM Ipomoeassin F (Ipom-F), a small molecule inhibitor of the Sec61 channel, (Zong et al., 2019), or a solvent control. Isolated membranes (B) and total translation reactions (C) were resolved by SDS-PAGE and results were visualised by phosphorimaging. Vpu, Sec61β and Ii were used as control membrane proteins that are insensitive (Vpu and Sec61β) or sensitive (Ii) to Ipom-F (O’Keefe et al., 2021b; O’Keefe et al., 2021c). Red dots indicate N-glycosylated protein species.

When LD proteins insert into the ER membrane with their N-terminal extension located inside the ER lumen they assume, albeit transiently, the same orientation as a stable type III transmembrane protein such as the Vpu protein (cf. Fig. 5A; see also (McKenna et al., 2017; O’Keefe et al., 2021b; O’Keefe et al., 2021c)). Since *bona fide* type III membrane proteins are integrated via a novel pathway that utilises the membrane insertase activity of the EMC (Chitwood et al., 2018; O’Keefe et al., 2021c), we explored the possibility that some LD membrane proteins may also utilise the EMC during their membrane insertion. To this end, we carried out *in vitro* translation reactions in the presence of semi-permeabilised (SP) cells depleted of specific ER components implicated in membrane protein biogenesis (cf. (Lang et al., 2012; O’Keefe et al., 2021b; O’Keefe et al., 2021c)). Briefly, HeLa cells transfected with siRNA oligonucleotides targeting specific membrane components were treated with digitonin to selectively permeabilise their plasma membrane, the cytosol was washed away and the resulting SP cells used as a source of membranes for *in vitro* translation reactions (cf. (Wilson et al., 1995)). On the basis of our present findings and those of previous studies (O’Keefe et al., 2021b; O’Keefe et al., 2021c; Schrul and Kopito, 2016) we chose to deplete: SRα, a subunit of the SRP receptor which enables the co-translational delivery of nascent precursor proteins to the ER membrane (O’Keefe et al., 2021a); EMC5, a core structural subunit of the EMC (Bai et al., 2020; Miller-Vedam et al., 2020; O’Donnell et al., 2020; Pleiner et al., 2020; Volkmar et al., 2019); Sec61α, a central component of the Sec61 translocon (O’Keefe et al., 2021a) and Pex3, a membrane-tethered component of the post-translational pathway used by UBXD8 (Schrul and Kopito, 2016).

Having confirmed ∼85-96% depletion of each of these membrane-localised factors (Fig. 6A and Table S1), we then tested how their loss affects the ER insertion of OPG2-tagged LD membrane proteins by comparing the amount of N-glycosylated products that were recovered relative to control SP cells (cf. Fig. S6A). Perturbation of the EMC dramatically reduced the amount of N-glycosylated, and hence fully membrane-spanning, polypeptides for all three co-translationally inserting LD membrane proteins tested (Fig. 6B, cf. 1g species in lanes 1 and 4 for OPG2-HSD17B11-FLAG, OPG2-METTL7B-FLAG and OPG2-AUP1-FLAG). In contrast, the insertion of Ii, a Sec61-dependent type II membrane protein (cf. Fig. 5A), was barely altered (Fig. 6B, cf. 2g species in lanes 1 and 4 for Ii). Quantification showed that EMC5 depletion reduced the integration of the three model LD membrane proteins that employ the co-translational pathway by >50%, (see Fig. 6C), an outcome that is directly comparable to the defect in membrane insertion observed with OPG2-Vpu, a bone fide type III membrane protein recently shown to utilise the EMC (O’Keefe et al., 2021c) (cf. Fig. 5A). Likewise, depletion of the SRP receptor, which acts upstream of the EMC insertase during the co-translational insertion of type III proteins (O’Keefe et al., 2021c), also significantly reduced the membrane insertion of two LD proteins, OPG2-METTL7B-FLAG and OPG2-AUP1-FLAG, the type III membrane protein OPG2-Vpu and the classical co-translational Sec61 client Ii (Fig. 6C). In contrast, although the membrane insertion of Sec61βOPG2, a post-translational tail-anchored protein client of the EMC (Guna et al., 2018) showed the strongest reduction of any protein tested following EMC5 depletion, loss of SRα had no significant effect *in vitro* (Fig. 6C). Hence, in contrast to such tail-anchored proteins, LD protein clients that employ the EMC co-translationally, also require the actions of the SRP receptor as previously reported for *bona fide* type III membrane proteins (O’Keefe et al., 2021c).

**Fig. 6.**
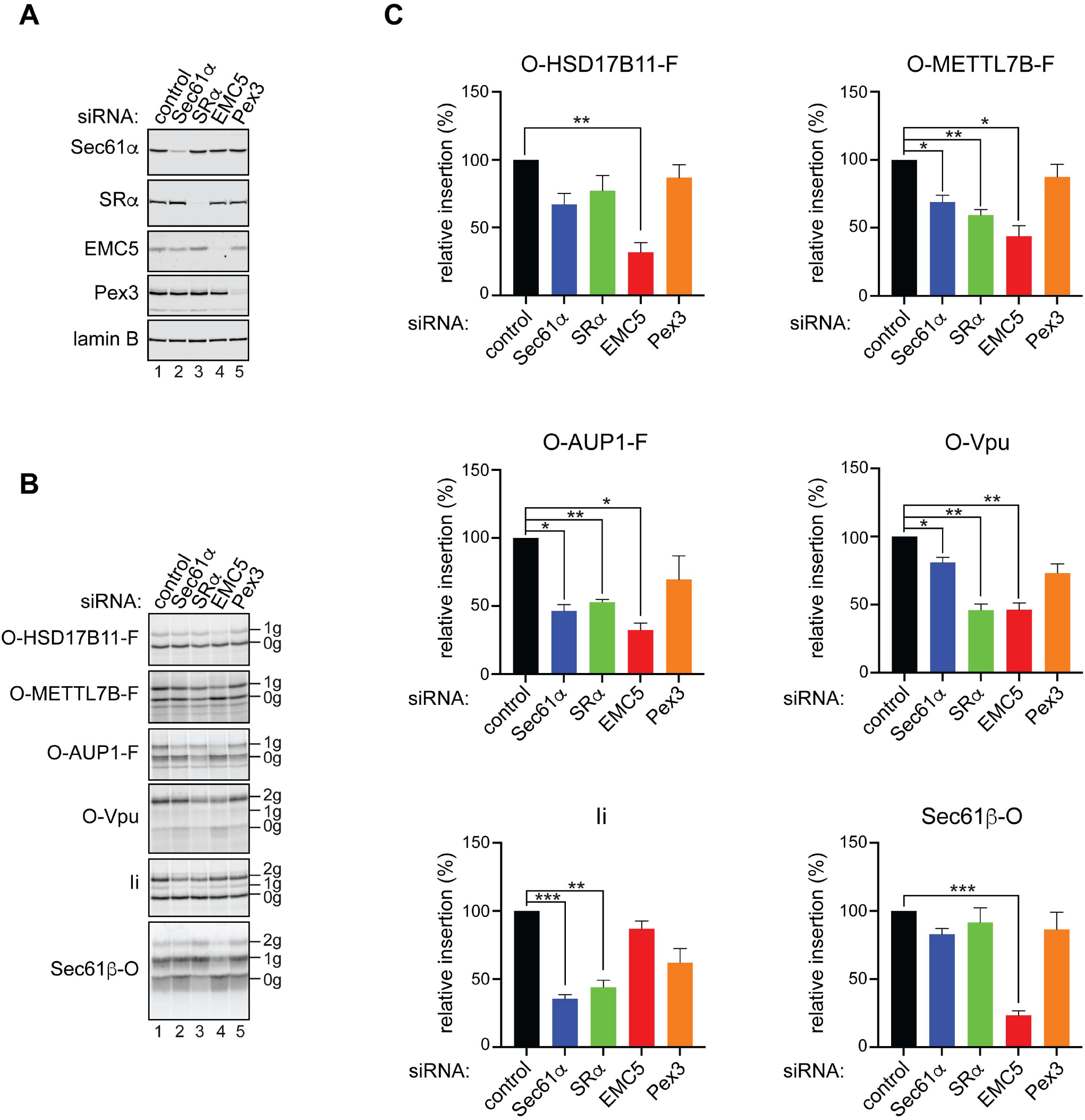
The EMC facilitates the biogenesis of LD membrane proteins at the ER. **(A)** HeLa cells were depleted of selected factors implicated in membrane protein biogenesis at the ER via siRNA-mediated knock-downs, their plasma membrane was then selectively permeabilised and the knock-down efficiency in these semi-permeabilised cells confirmed by Western blotting with antibodies against the indicated proteins (see also Table S1). **(B)** The semi-permeabilised HeLa cells (see panel A) were used as a source of ER membrane during *in vitro* synthesis of the indicated LD membrane proteins tagged with the OPG2 epitope (“O”) at the N-terminus and the FLAG tag (“F”) at the C-terminus. In parallel, OPG2-Vpu, an EMC-dependent type III membrane protein (O’Keefe et al., 2021b; O’Keefe et al., 2021c), the tail-anchored protein Sec61β-OPG2 and untagged invariant chain (Ii), a classical Sec61-dependent substrate, were synthesised. Membrane fractions were isolated, resolved by SDS-PAGE and results visualised by phosphorimaging. “0g” indicates non-glycosylated, “1g” singly N-glycosylated and “2g” doubly N-glycosylated protein species. **(C)** ER membrane insertion efficiency, defined as the amount of N-glycosylated species in the membrane fraction and normalised to values obtained for the non-targeting siRNA, was calculated for each of the proteins and knock-down conditions. Shown are the means with error bars indicating standard error of means. Statistical significance was calculated using repeated measures (RM) one-way ANOVA for n ≥ 3 biological replicates. * p ≤ 0.05, ** p ≤ 0.01, *** p ≤ 0.001.

Interestingly, although the membrane insertion of LD membrane proteins via this co-translational pathway is insensitive to the Sec61 inhibitor, Ipom-F (Fig. 5), two of these proteins, OPG2-METTL7B-FLAG and OPG2-AUP1-FLAG, showed a significant loss of membrane insertion following Sec61α depletion (Fig. 6C). This mirrors the recently reported behaviour of type III membrane proteins ((O’Keefe et al., 2021c); see also Fig 6C, OPG2-Vpu) and suggests that the Sec61 translocon may contribute to LD membrane protein biogenesis via a mechanism that does not require its insertase activity (see Discussion). Despite the effectiveness of its knockdown (∼90%, see Table S1), Pex3 depletion had no effect on the membrane insertion of any of the proteins tested (Fig. 6C), consistent with its role as a receptor for the post-translational insertion of LD proteins with pre-formed hairpin loops into the cytosolic leaflet of the ER membrane (Schrul and Kopito, 2016). In summary, our studies suggest that a subset of LD membrane proteins is synthesised via an alternative, co-translational, pathway that initially involves their EMC-mediated insertion as “type III-like” membrane proteins. Unlike *bona fide* type III membrane proteins, these LD membrane proteins are subsequently able to dislocate their short hydrophilic N-termini into the cytosol, thereby generating a stable hairpin loop conformation that is competent for incorporation into newly forming LDs.

## DISCUSSION

LD membrane proteins regulate key aspects of LD function and dynamics (Olzmann et al., 2013; Wilfling et al., 2013; Zhang et al., 2017), and have been directly linked to cancer (VandeKopple et al., 2019; Zhang et al., 2017) and viral propagation (Park et al., 2015; Zhang et al., 2018). They initially insert into the ER before being trafficked to LDs (Ingelmo-Torres et al., 2009; Ostermeyer et al., 2004; Schrul and Kopito, 2016; Stevanovic and Thiele, 2013; Turro et al., 2006; Zehmer et al., 2009; Zehmer et al., 2008). Early studies using heterologous systems suggested that plant oleosins, authentic LD membrane proteins, and mammalian caveolins, which localise to LDs under specific conditions, associate with the ER membrane co-translationally in a process mediated by SRP, SR and, for caveolins, the Sec61 translocon (Abell et al., 2002; Beaudoin et al., 2000; Monier et al., 1995). However, a more recent report (Schrul and Kopito, 2016) established that the model LD membrane protein, UBXD8, is delivered to the ER post-translationally via a pathway that is mediated by Pex19 and its ER membrane-localised receptor, Pex3. This Pex19/Pex3-dependent pathway is also suggested to accommodate other hairpin-forming, ER-resident proteins (Yamamoto and Sakisaka, 2018) and hence was proposed as a general pathway for the ER membrane insertion of LD-destined membrane proteins (Dhiman et al., 2020; Schrul and Kopito, 2016; Yamamoto and Sakisaka, 2018).

Here, we have used a homologous mammalian cell-free translation system together with cell-based studies to investigate the biogenesis of a panel of well-defined human LD membrane proteins at the ER. We find that LD membrane proteins can be divided into at least two different groups, which show distinct requirements for delivery to and insertion into the ER (Fig. 7). The first group is exemplified by UBXD8 and HSD17B7, and follows a post-translational targeting pathway as previously defined (Schrul and Kopito, 2016) (cf. Fig. 1B). The second group relies on a co-translational route for their efficient delivery to the ER, and its clients include AUP1, HSD17B11, METTL7A, METTL7B and HIG2 (cf. Fig. 1B). The biogenesis of these co-translationally inserted LD membrane proteins is mediated by the EMC, with additional contributions from SR and the Sec61 translocon detected for METTL7B and AUP1 (cf. Fig. 6). The ER insertion of such LD membrane proteins is consistently insensitive to Ipom-F (cf. Fig. 5), a small molecule inhibitor of the Sec61 translocon (Zong et al., 2019), arguing that it is not acting as an insertase in this context. Rather, we speculate that the Sec61 complex can enhance the co-translational insertase activity of the EMC as recently established for the biogenesis of *bona fide* type III membrane proteins (cf. (O’Keefe et al., 2021c)).

**Fig. 7.**
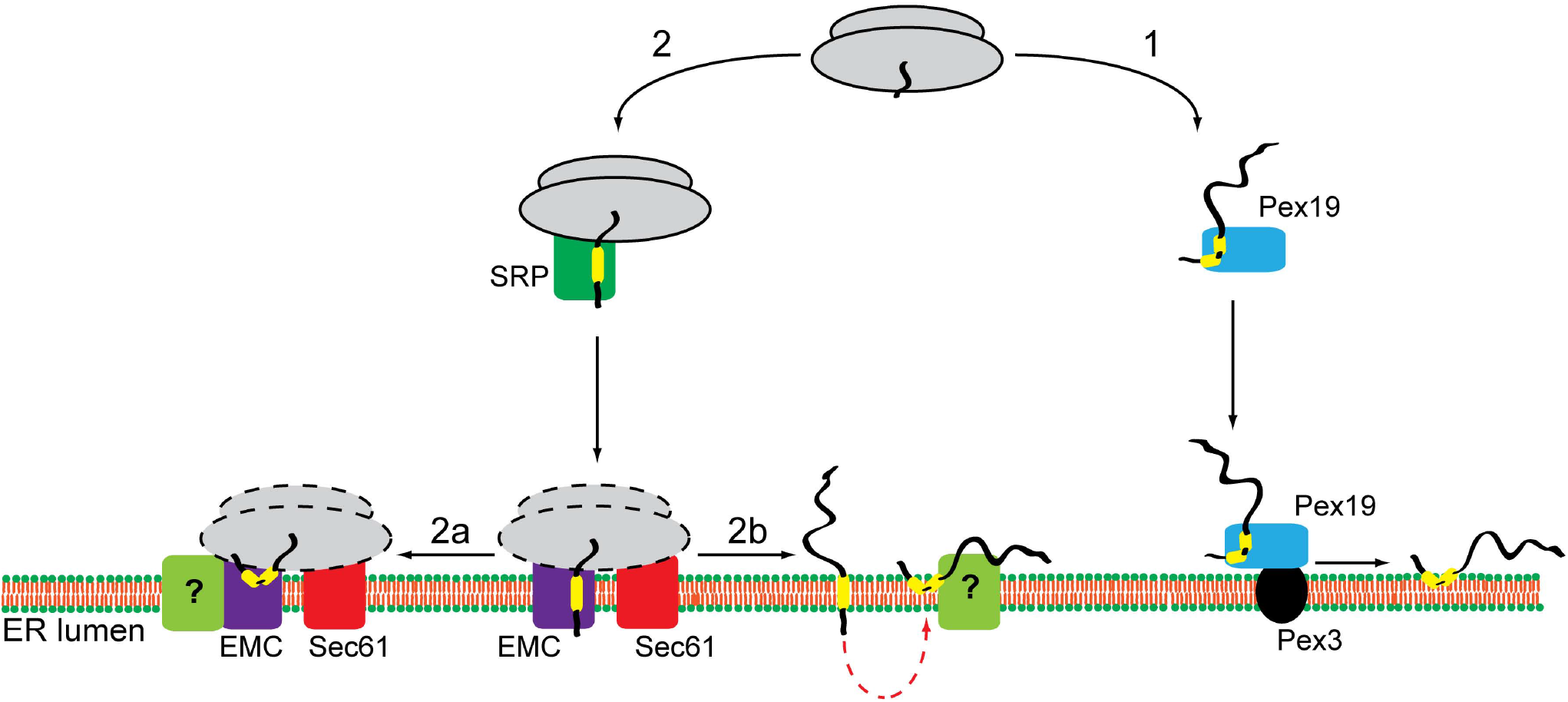
Model for LD membrane protein biogenesis at the ER. Depending on the properties of their hydrophobic, membrane-inserting region, LD membrane proteins enter one of two potential biosynthetic pathways. Proteins such as UBXD8 and HSD17B7 are released into the cytosol, bound by Pex19 and post-translationally delivered to the ER (route 1). Interaction between Pex19 and its membrane-tethered receptor, Pex3, releases such LD membrane proteins in a hairpin conformation into the ER membrane. Alternatively, LD membrane proteins such as HSD17B11, METTL7A, METTL7B, AUP1 and HIG2, are delivered co-translationally to the ER and insert into the lipid bilayer in a reaction facilitated by the EMC with additional, insertase-independent, contributions from the Sec61 complex (route 2). At the moment, it is unclear how the EMC and the Sec61 complex co-operate with each other, and which of these two complexes provides a binding site for the ribosome (indicated by dashed lines). The co-translationally delivered LD membrane proteins transiently expose their N-terminus to the ER lumen but then re-arrange their topology to form a hairpin with both termini facing the cytosol, which is a prerequisite for trafficking to LDs. This topological re-orientation could occur either co-translationally (route 2a) or following complete protein synthesis (route 2b). At present, it is unknown whether this process is spontaneous or requires an assistance from a dedicated factor(s) (indicated with “?”).

Our data suggest that the EMC may not discriminate between proteins that stably reside in a membrane-spanning topology and proteins that can subsequently acquire a hairpin conformation, consistent with the ability of the EMC to translocate both N-terminal and C-terminal segments of hydrophilic polypeptide that are adjacent to a hydrophobic transmembrane region (Bai et al., 2020; Chitwood et al., 2018; Guna et al., 2018; Miller-Vedam et al., 2020; O’Donnell et al., 2020; Pleiner et al., 2020). Our finding that co-translationally inserted LD membrane proteins can typically expose their short N-termini to the ER lumen (cf. Fig. 2) confirms previous studies using chimeric LD proteins with artificial N-terminal signal sequences. Hence, signal sequence bearing versions of METTL7A (Zehmer et al., 2009) and AUP1 (Stevanovic and Thiele, 2013) are efficiently targeted to the ER, processed by ER lumenal signal peptidase (Paetzel et al., 1998), and the resulting signal sequence-cleaved proteins incorporated into LDs (Stevanovic and Thiele, 2013; Zehmer et al., 2009). Likewise, in this study we show that N-glycosylated HSD17B11 can reach mature LDs (cf. Figs 3A and 3B). Hence, exposure of the N-termini of such LD proteins to the ER lumen, whether by virtue of an artificial signal sequence (Stevanovic and Thiele, 2013; Zehmer et al., 2009) or not (this study), does not prohibit either their acquisition of a hairpin topology or their authentic LD localisation. Taken together, these studies strongly support the idea that the co-translational biogenesis of LD membrane proteins at the ER generates a functional pool of polypeptides that can be trafficked to LDs rather than an “off-target” pathway.

An obvious consequence of our model for the EMC-mediated co-translational biogenesis of LD proteins is that at some point they must reorient from a fully membrane-spanning topology to a hairpin one, in order to be accommodated by LDs (cf. (Abell et al., 2002; Stevanovic and Thiele, 2013; Zehmer et al., 2009; Zehmer et al., 2008)). The normally transient nature of their N-terminal domain residency in the ER lumen is also underlined by our protease protection studies of LD membrane proteins. Hence, we only detect membrane-protected, N-terminal fragments of these proteins in the presence of N-linked glycans (cf. Figs 3C and S5), which most likely “trap” these otherwise labile topological intermediates (Goder et al., 1999). At present we can only speculate as to how the re-orientation of such LD membrane proteins can occur. One possibility is that the TMD acquires its hairpin conformation co-translationally (Fig. 7, route 2a). This would likely result in the brief exposure of the N-terminus to the ER lumen, consistent with the incomplete N-glycosylation of most LD-membrane proteins that we observe (cf. Fig. 2). In this scenario, LD membrane protein biogenesis would resemble that of some type II membrane proteins which initially insert “head-first” (N-terminus translocated) and subsequently completely invert their topology within the Sec61 translocon so that their N-terminus faces the cytosol (Devaraneni et al., 2011; Goder and Spiess, 2003). Alternatively, LD membrane proteins might complete their synthesis, be released into the ER membrane and only then reorient their topology (Fig. 7, route 2b) as reported for some bacterial membrane proteins in response to changes in the membrane lipid composition (Dowhan et al., 2019).

How LD membrane proteins can transition from a fully membrane-spanning topology to a hairpin one and whether this process is spontaneous or requires dedicated factors will be key questions for future studies. Notably, when the hairpin motif of a LD membrane protein is replaced with a “classical” TMD the resulting polypeptide does not localise to LDs (Zehmer et al., 2008). Furthermore, the ER-LD interface appears to act as a barrier that can exclude fully membrane-spanning proteins from LDs (Khaddaj et al., 2022). Hence, it seems that specific characteristics of the membrane-embedded region of LD membrane proteins are likely to be responsible for their ability to rearrange into a hairpin. Proline residues, which act as α-helix breakers (MacArthur and Thornton, 1991), are often found within the hairpin motif of LD membrane proteins and might possibly constitute one such determinant for reorientation. Indeed, a Pro-Val-Gly motif is needed for AUP1 to achieve a hairpin topology (Stevanovic and Thiele, 2013), whilst the so-called proline-knot motif is required for oleosin trafficking to LDs, although not for their insertion into the ER membrane (Abell et al., 1997). At the same time, mutagenesis of proline residues in METTL7A, METTL7B, Cyb5R3 and caveolins (Ostermeyer et al., 2004; Zehmer et al., 2008) does not affect their ER insertion and LD trafficking, suggesting a more complex relationship between protein sequence and membrane topology.

Such complexity is further underpinned by our TMD swap experiments (cf. Fig. 4) where we find that the TMD of METTL7B can support the translocation of ∼90 N-terminal residues of UBXD8 into the ER lumen, whilst when UBXD8 hairpin is incorporated into METTL7B it almost completely abolishes the otherwise efficient translocation of its much shorter N-terminus. It seems likely that in addition to features such as hydrophobicity, the folding of the TMD/hairpin loop region may also influence which biosynthetic route is taken, for example, by impacting on the recruitment of targeting factors or the ability of a particular client to engage the EMC. Whilst our study indicates that the TMD/hairpin motif is the primary determinant for pathway selection it is possible that specific sequence features of its flanking regions can influence the efficiency of pathway entry. For example, replacing the ∼100 residues located N-terminally to the TMD/hairpin of caveolin-1 with a region of a comparable length derived from rat growth hormone changes the topology of a fraction of the protein from a hairpin loop to a fully membrane-spanning (Monier et al., 1995). It should also be noted that our studies of RHD14 indicate that at least some LD proteins can utilise both routes effectively, suggesting that these pathways are not mutually exclusive, but rather that some clients are more dependent on one or other as also reported for the biogenesis of tail-anchored proteins at the ER (cf. (Casson et al., 2017)).

In summary, we show that LD membrane proteins can be delivered to the ER via at least two distinct pathways, that can most simply be classified as co-translational (this study) and post-translational (Schrul and Kopito, 2016). Our work reconciles some apparent discrepancies between previous publications (Abell et al., 2002; Beaudoin et al., 2000; Monier et al., 1995; Schrul and Kopito, 2016) and lays the foundations for future studies aimed at unravelling the complexity of LD membrane protein biogenesis that we have revealed.

## MATERIALS AND METHODS

### Materials

LD membrane protein cDNAs in pcDNA3.1+/C-(k)DYK were purchased from GenScript and their variants with the OPG2 tag, AUP1 point mutants and METTL7B/UBXD8 chimaeras were generated by site-directed mutagenesis. Sec61β with a C-terminal OPG2 tag in pcDNA5, either with or without an N-terminal FLAG epitope, was previously described (Leznicki and High, 2020). Nuclease-treated rabbit reticulocyte lysate was from Promega (L4960), [^35^S] methionine (EasyTag EXPRESS ^35^S Protein Labelling Mix) from Perkin Elmer, LipidToxRed stain (H34476) from ThermoFisher Scientific and Endoglycosidase H (P0702 and P0703) from New England Biolabs. The hybridoma line producing the monoclonal anti-rhodopsin antibody (Adamus et al., 1991) was provided by Paul Hargrave (Department of Ophthalmology, University of Florida, US), monoclonal anti-tubulin antibody by Keith Gull (University of Oxford, UK), rabbit polyclonal anti-Sec61α antibody was a gift from Richard Zimmermann and Sven Lang (Saarland University, Homburg, Germany) whilst rabbit anti-SRα antibodies (Jadhav et al., 2015) were gifted by Martin Pool (University of Manchester, UK). The following commercial antibodies were used: rabbit anti-β-actin (Abcam, cat. number ab8227, 1:5000, batch numbers GR3176830-1 and GR3224338-1), mouse anti-FLAG (clone M2, Sigma Aldrich, cat. number F3165, 1:2500, batch number SLBH1191V), rabbit anti-FLAG (Sigma Aldrich, cat. number F7425, 1:1000), anti-AUP1 (Bethyl Laboratories, cat. number A302-899A, 1:1000, batch number 1), anti-BAP31 (ProteinTech, cat. number 11200-1-AP, 1:1000, batch number 00018537), anti-ADRP (Abcam, cat. number ab78920, 1:1000, batch number GR3227109-1), anti-calnexin (Cell Signaling Technology, clone C5C9, cat. number 2679S, 1:2000, batch number 4), anti-MMGT1 (EMC5) (Bethyl Laboratories, cat. number A305-832A-M, 1:1000, batch number 1), anti-Pex3 (St John’s Laboratory, cat. number STJ29491, 1:1000, batch number 949135460201), anti-lamin B (SantaCruz Biotechnology, cat. number sc-6217, 1:1000, batch number J1311). MMGT1 (EMC5) siRNA was obtained from ThermoFisher Scientific (cat. number s41129) whilst all the other siRNAs were made to order as “on-target+” by Horizon Discovery. U2OS cells were procured from ECACC whilst HeLa and HepG2 cells were from ATCC, and all were checked for mycoplasma infection. Ipomoeassin F was synthesised as previously described (Zong et al., 2015; Zong et al., 2020; Zong et al., 2017).

### *In vitro* transcription and translation

Templates for *in vitro* transcription were generated by PCR, mRNA was prepared as previously described (Leznicki et al., 2013) and used in *in vitro* translation reactions comprising rabbit reticulocyte lysate, 1 mCi/ml [^35^S] methionine and amino acid mix lacking methionine. Where indicated, reactions were also supplemented with 1 μM Ipomoeassin F or an equal volume of DMSO solvent. To compare co- and post-translational insertion into the ER, LD membrane proteins and control proteins were synthesised in parallel either in the presence or absence of ER-derived dog pancreatic microsomes at 30°C for 7 min and further translation initiation was blocked with 0.1 mM aurintricarboxylic acid (Alfa Aesar, cat. number A15905). Samples were incubated at 30°C for a further 8 min, puromycin (Sigma-Aldrich, cat. number 540222) added to a final concentration of 2.5 mM and reactions kept at 30°C for 7 min. At this stage the “post-translational” reactions were supplemented with ER-derived microsomes whilst the “co-translational” reactions with an equal volume of KHM buffer (110 mM KOAc, 2 mM Mg(OAc)_2_, 20 mM HEPES-KOH, pH 7.5), and all samples were kept at 30°C for 15 min. Aliquots (∼7%) were taken to check total translation products whilst the remaining samples were spun through a high-salt sucrose cushion (0.75 M sucrose, 0.5 M KOAc, 5 mM Mg(OAc)_2_, 50 mM Hepes-KOH, pH 7.9) at 100,000 × g for 10 min at 4°C in order to isolate the membranes and membrane-associated material. The pellets were then directly resuspended in SDS sample buffer.

To check N-glycosylation of *in vitro* synthesised proteins, translations were carried out at 30°C for 15 min, further translation initiation was blocked with 0.1 mM aurintricarboxylic acid and samples were incubated at 30°C for 30 min. Total translation aliquots were taken, membranes isolated as described above, lysed directly in SDS samples buffer and proteins were denatured at 37°C for 30 min. Samples were then split in two and either buffer control or ∼20,000 U/ml of Endoglycosidase H variant added, followed by incubation at 37°C for at least 2h.

All samples were resolved by SDS-PAGE and results visualised by phosphorimaging using a Typhoon FLA 7000 phosphorimager (GE Healthcare). Images were processed and band intensity quantified using AIDA software (Raytek).

### Cell culture and preparation of semi-permeabilised (SP) cells

All cells were cultured in DMEM (Sigma-Aldrich, D5796) supplemented with 10% (v/v) foetal bovine serum and were maintained in a 5% CO_2_ humidified incubator at 37°C. Transient transfection of U2OS and HepG2 cells with plasmid DNA was carried out using GeneJuice (Merck Millipore, cat. number 70967) according to manufacturer’s instruction and keeping GeneJuice to DNA ratio of 3:1 (volume in μl : weight in μg). To estimate the N-glycosylation of the proteins studied, U2OS cells grown in 6-well plates were transfected with 2 μg of plasmid DNA and lysed directly in SDS sample buffer 24h post-transfection, whilst the amount of DNA used for other experiments is described in detail in the relevant sections of Materials and Methods (see below). Depletion of membrane components was carried out in HeLa cells using INTERFERin (Polyplus, cat. number 409-10) as a transfection reagent according to manufacturer’s instruction. The following siRNA oligonucleotides were used at 20 nM final concentration: non-targeting siRNA (sequence UGGUUUACAUGUUGUGUGAuu), SEC61A1 (Sec61α) siRNA (sequence AACACUGAAAUGUCUACGUUUuu), SRPRA (SRα) siRNA (sequence GAGCUUGAGUCGUGAAGACuu), PEX3 siRNA (sequence GGGAGGAUCUGAAGAUAAUAAGUUUuu) and MMGT1 (EMC5) siRNA (ThermoFisher Scientific, s41129). 850,000 cells were plated in a 10-cm dish, transfected with siRNA oligonucleotides the next day and grown for another 72h, at which point SP cells were prepared. To this end, cells were harvested by trypsinisation in 3 ml 0.25% trypsin-EDTA solution (Sigma-Aldrich, T3924), which was then inhibited by adding 4 ml of ice-cold KHM buffer (110 mM KOAc, 2 mM Mg(OAc)_2_, 20 mM HEPES-KOH, pH 7.5) supplemented with 100 μg/mL Soybean trypsin inhibitor (Sigma-Aldrich, T6522). Cells were pelleted at 500 × g for 3 min at 4°C, resuspended in 4 ml ice-cold KHM buffer supplemented with 80 μg/ml high purity digitonin (Calbiochem, cat. number 300410) and incubated on ice for 5 min to permeabilise the plasma membrane. Cells were diluted to 14 ml with ice-cold KHM buffer, pelleted at 500 × g for 3 min at 4°C, resuspended in 5 ml ice-cold HEPES buffer (90 mM HEPES-KOH, pH 7.5, 50 mM KOAc) and incubated on ice for 10 min. Cells were pelleted by centrifugation once more, resuspended in 100 ul KHM buffer and endogenous mRNA removed by treatment with 0.2 U Nuclease S7 Micrococcal nuclease from *Staphylococcus aureus* (Sigma-Aldrich, 10107921001) in the presence of 1 mM CaCl_2_ at 22°C for 12 min. Nuclease was inactivated by the addition of EGTA to 4 mM final concentration, SP cells centrifuged at 13,000 × g for 1 min and resuspended in KHM buffer to a final concentration of 3×10^7^ cells/ml. *In vitro* translation reactions were carried out for the indicated proteins using rabbit reticulocyte lysate and SP cells at a final concentration of 3×10^6^ cells/ml at 30°C for 40 min in a thermomixer (Eppendorf) set to 900 rpm. Reactions were placed on ice, 10% (v/v) kept as an input material, the rest diluted with 1 ml ice-cold KHM buffer and the SP cells isolated by centrifugation (21,000 × g, 2 min, 4°C) followed by lysis in SDS sample buffer.

### Isolation of LDs

LDs were isolated based on the protocol from (Ingelmo-Torres et al., 2009) with minor modifications. HepG2 cells in a 15-cm dish were transfected at ∼50% confluency with 10 μg of pcDNA3.1+/C-(k)DYK plasmid encoding OPG2-HSD17B11-FLAG. 24h post-transfection LD formation was induced by adding fresh medium supplemented with 0.5 mM oleic acid (Sigma Aldrich, O1383) complexed with bovine serum albumin (Sigma Aldrich, A8806) (Brasaemle and Wolins, 2016), and zVAD-fmk (Selleck Chemicals, S7023) was added to a final concentration of 50 μM to inhibit N-glycanase (Misaghi et al., 2004). After 16h cells were washed twice with ice-cold PBS, harvested by scraping, pelleted (500 × g, 5 min, 4°C) and resuspended in 0.5 ml buffer L (50 mM Tris-Cl, pH 7.5, 150 mM NaCl, 5 mM EDTA) supplemented with a complete protease inhibitor cocktail (Sigma Aldrich, P8340). Cells were broken by passing 30 times through a cell homogeniser (Isobiotec, Germany) with a tungsten carbide ball of 14 μm clearance. Cell lysate was pre-cleared (1,600 × g, 5 min, 4°C), 0.5 ml of supernatant mixed with 0.5 ml of 2.5M sucrose in buffer L, and overlaid with 200 μl of 30%, 25%, 20%, 15%, 10%, 5% (w/v) sucrose in buffer L. Samples were centrifuged at 50,000 rpm (∼166,000 × g) for 3h at 4°C using TLS-55 rotor (Beckman Coulter) in an Optima benchtop ultracentrifuge (Beckman Coulter) with acceleration set to 9 and deceleration set to 0. Five fractions (280 μl each) were collected from the top using a Hamilton syringe followed by the final ∼700 μl bottom fraction, and an aliquot of each fraction was mixed directly with SDS sample buffer and resolved by SDS-PAGE for Western blotting analysis.

To quantify N-glycosylated OPG2-HSD17B11-FLAG in the LD fraction that is authentically LD-localised (*Ngly*. 17*B*11_*LD specific*_) or ER-associated (*Ngly*. 17*B*11_*ER contaminants*_) the following formulas were used:

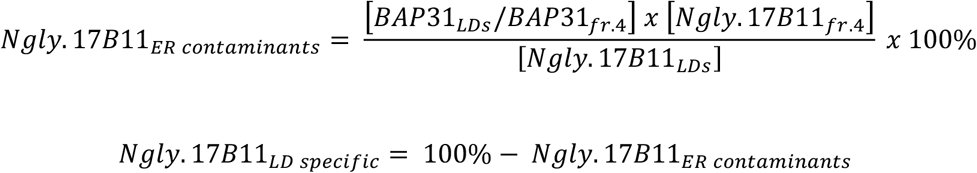

where [*Ngly*.. 17*B*11_*LDs*_] corresponds to total N-glycosylated OPG2-HSD17B11-FLAG in the LD fraction, [*Ngly*.. 17*B*11_*fr.*4_ to total N-glycosylated OPG2-HSD17B11-FLAG in fraction 4 (main ER fraction) and *BAP*31_*LDs*_/ *BAP*31_*fr*.4_] is the ratio between BAP31 levels in the LD fraction and fraction 4.

### Protease protection assay

HeLa cells grown in 10-cm dishes were transfected with 5 μg of the indicated LD membrane protein variants in pcDNA3.1+/C-(k)DYK or Sec61β in pcDNA5, and 24h post-transfection SP cells were prepared as described above (“Cell culture and preparation of semi-permeabilised (SP) cells” section) but omitting the nuclease treatment step. SP cells were resuspended to a final concentration of ∼2.5×10^7^ cells/ml and split into three aliquots, which received: water, 1 mg/ml proteinase K (Sigma Aldrich, P2308) only or 1 mg/ml proteinase K together with 1% (w/v) Triton X-100. Reactions were incubated for 1h at 22°C in a thermomixer (Eppendorf) with mixing set to 1000 rpm, protease was inhibited with 2.5 mM PMSF and samples incubated for another 10 min (22°C, 1000 rpm). Reactions were stopped by adding hot SDS sample buffer and immediate incubation at 95°C for 10 min. To reduce the viscosity of samples, DNA was sheared using a BioRuptor (Diagenode).

### Fluorescence microscopy

U2OS cells were grown on glass coverslips in 6-well plates and at ∼40% confluency transfected with 1 μg of plasmids encoding the indicated proteins using GeneJuice as described above. Media were replaced ∼6h post-transfection with fresh DMEM supplemented with 0.25 mM oleic acid complexed with bovine serum albumin (Brasaemle and Wolins, 2016), and the cells were grown for another 16h. Cells were fixed with 4% (w/v) paraformaldehyde for 15 min at RT, washed 3 times with PBS supplemented with 100 mM glycine pH 8.0 (5 min each wash step) and finally washed with PBS without glycine. At this stage their plasma membrane was permeabilised with 20 μM digitonin in PBS for 5 min at RT, coverslips were washed 3 times with PBS, blocked with 1% (w/v) bovine serum albumin in PBS for 15 min at RT and washed once again with PBS. Coverslips were then incubated with the anti-FLAG antibody (clone M2, 1:800 dilution in 1% (w/v) bovine serum albumin in PBS) for 1h at RT, washed 3 times with PBS (5 min each wash step) and incubated for 1h at RT with a secondary donkey anti-mouse antibody conjugated to AlexaFluor488 dye (1:1000 dilution in 1% (w/v) bovine serum albumin in PBS). After washing 3 times with PBS (5 min each wash step) the coverslips were incubated with LipidToxRed dye (1:400 in PBS) for 1h at RT, briefly washed and mounted using ProLong Gold (ThermoFisher Scientific, P36930). Images were acquired on an Olympus IX83 inverted microscope using 60x/1.42 Plan Apo objective and a CCD camera with a Z optical spacing of 0.2 μm. Raw image deconvolution was carried out using Huygens Pro software (SVI) and images were processed with ImageJ (Fiji).

### Statistical analysis

Radiolabelled protein species were quantified using AIDA software whilst signals resulting from Western blotting were quantified with ImageStudioLite (LiCor). Calculations were carried out in Microsoft Excel and GraphPad Prism was then used to generate graphs and quantify statistical significance using the indicated tests.

## ACKNOWLEDGEMENTS

For gifting the reagents, we thank Richard Zimmermann and Sven Lang (anti-Sec61α antibody; University of Saarland, Homburg, Germany), Martin Pool (anti-SRα antibody, University of Manchester, UK) and Sarah O’Keefe (AUP1^E2N/P4T^ cDNA, University of Manchester, UK). We are also grateful to Martin Lowe, Martin Pool, Sarah O’Keefe and Lisa Swanton (University of Manchester, UK) for critical feedback during the preparation of the manuscript.

